# Cardiometabolic state links neurovascular burden with brain structure and function across age: evidence from EEG and MRI

**DOI:** 10.1101/2024.05.31.596817

**Authors:** Daniel Franco-O’Byrne, Ana M. Castro-Laguardia, Carolina Delgado, James M. Shine, David Huepe, Enzo Tagliazucchi, Cecilia Gonzalez Campo, Agustín Ibañez, Vicente Medel

## Abstract

Aging affects brain structure and function alongside metabolic and vascular processes leading to energetic impairments. While local neurometabolic dysfunction in aging is well-documented, the influence of systemic cardiometabolic and vascular markers on brain structure and function remains less understood. We examine the link between cardiometabolic dysfunction (measured by an allostatic load index) and neurovascular burden (measured by white matter hyperintensities) with brain changes, including ventricular and hippocampal volume, as well as EEG activity, across age. Analyzing data from 196 healthy individuals across age (20-75 years), we found a significant positive correlation between allostatic load index and white-matter hyperintensities, irrespective of age. White-matter hyperintensities are also positively linked with ventricular enlargement, but not hippocampal atrophy. The allostatic load index mediated the relationship between white-matter hyperintensities and ventricular volume. Regarding brain function, changes in the spectral aperiodic exponent but not periodic alpha power were linked to white-matter hyperintensities and the allostatic load index. Such index mediated the relationship between spectral aperiodic exponent and white-matter hyperintensities. Thus, findings suggest that the cardiometabolic state, as measured by an allostatic load index, plays a crucial role in brain health across age, particularly influencing ventricular enlargement and increased aperiodic activity.

## Introduction

The human brain involves dynamic lifespan crosstalks between neural activity and multisystem metabolic processes (Li & Sheng, 2022). These energetic requirements are met by direct supply through specialized neurometabolic pathways (Li & Sheng, 2022; Figley & Stroman, 2011; García-Cazorla & Saudubray, 2018; Thakur et al., 2022) and global cardiometabolic processes fueling bodily-level organs, including the brain (Pontzer et al., 2016) where both metabolic modalities are affected across aging (Ainslie & Bailey, 2013; Cunnane et al., 2020; Gruenewald et al., 2009; Shiels et al., 2017). Specifically, brain structure -brain gray matter and ventricular volume-, and brain function -neural activity of EEG dynamics- are both constrained by age-related metabolic dysfunction (Goyal et al., 2017; Ibáñez et al., 1998; Moretti et al., 2017; Shah et al., 2017). While research has identified explicit mechanistic pathways linking neural structure and function to neurometabolic supply (Li & Sheng, 2022; Medel et al., 2022), understanding the impact of body-level cardiometabolic status on brain health remains elusive.

White matter hyperintensities (WMHs) seen in T2 sequences of brain MRI reflect white matter lesions traditionally considered of vascular origin (Claassen et al., 2021). These are commonly found in middle-aged and elderly individuals. While WMHs have been linked to small vessel disease (Goyal et al., 2017; Ibáñez et al., 1998; Lambert et al., 2016; Moretti et al., 2017; Shah et al., 2017), they have also been related to inflammation (Nam et al., 2022; Raz et al., 2012), neurodegeneration (Dadar et al., 2020; Rizvi et al., 2021), and disruptions in brain dynamics (Atwi et al., 2018; Babiloni et al., 2008; Kumral et al., 2022). Additionally, evidence has shown that WMHs are also associated with both local neurometabolic processes (Brier et al., 2022; Jiaerken et al., 2019) as well as global systemic cardiometabolic dysregulation (Lampe et al., 2019). Thus, WMHs represent a structural biomarker of neurovascular burden, offering a window to integrate systemic cardiometabolic pathways with brain structure and function.

The systemic cardiometabolic state can be effectively operationalized by allostasis (Fava et al., 2019; Franco-O’Byrne et al., 2024.; Guidi et al., 2021). Allostasis refers to anticipatory regulation of bodily, cardiac, metabolic, and inflammatory processes to maintain the organism’s stability and adaptation (Schulkin & Sterling, 2019; Sterling, 2012). The allostatic load index (ALI), which integrates these measures in a single score, has been related to diverse functional and structural changes in the brain leading to inflammatory, neurovascular, and neurodegenerative statuses (Butterfield et al., 2022; de la Monte, 2017; Furlan & Petrus, 2023; Kapogiannis, 2015; López-Ojeda & Hurley, 2023; Roh et al., 2016). Indeed, chronic allostatic dysregulation, as indicated by a high ALI, triggers an exaggerated stress response within the brain, marked by the release of glucocorticoids, catecholamines, and proinflammatory cytokines leading to brain gray matter atrophy (Kline & Mega, 2020; Lenart-Bugla et al., 2022) as well as ventricular enlargement (McEwen, 2006, 2016). Consistently, allostatic overload-driven inflammatory states have been implicated in brain atrophy in hippocampal and prefrontal areas (Karoly et al., 2021; Lowther MK et al., 2020; Marsland et al., 2008; Schmidt MF et al., 2016) as well as increased WMHs (Nam et al., 2022; Raz et al., 2012), while also affecting brain electrophysiological dynamics (Düsing et al., 2016; Marshall & Cooper, 2017). Although the above evidence illustrates the associations between ALI and structural, neurovascular, and neurofunctional aspects, no consistent evidence of mechanisms linking these factors exists.

We addressed this gap by analyzing structural brain MRI, scalp EEG and cardiometabolic state to characterize the relationship of systemic metabolism with structural and functional brain signatures in aging. Our results consistently show a specific mediating mechanism of cardiometabolic state -as measured by ALI-between WMHs and functional brain activity, as well as with structural ventricular enlargement, but not hippocampal atrophy. These results suggest that multiple brain structural and functional signatures reflect an underlying cardiometabolic pathway across the lifespan.

## 2. Methods

### 2.1 Participants

We leverage the LEMON database data acquired in Leipzig (Germany) between 2013 and 2015 (https://fcon_1000.projects.nitrc.org/indi/retro/MPI_LEMON.html). Data was collected in accordance with the declaration of Helsinki and the study protocol was approved by the ethics committee at the medical faculty of the University of Leipzig (reference number 154/13-ff). The database is available under the Open Data Common License. The database originally consisted of 227 subjects. After converging subjects with MRI, EEG, and multisystemic data, 196 (69 females) subjects remained and were included in the present study’s analyses, with ages between 20 and 75 years and a mean education level of 12.45 ± 1.65 years. Table 1 shows a summary description of the sample.

**Table 1.** Sample demographic description.

| Table 1. Sample demographic description. |  |  |
| --- | --- | --- |
| Demographics |  | Total |
| Number of subjects |  | 196 |
| Years of age | (mean +- SD) | 38.77 (20.22) |
| Handedness | Right | 172 (87,8%) |
|  | Left | 20 (10,2%) |
|  | Ambidextrous | 4 (2%) |
| Education* | H | 4 (2%) |
|  | R | 37 (18,9%) |
|  | G | 153 (78,1%) |
|  | Not specified | 2 (1%) |

\*According to the German educational level's classification (G = Gymnasium (high school), R = Realschule (junior high school), H = Hauptschule (vocational school combined with apprenticeship)).

### 2.2. Data acquisition parameters

#### 2.2.1. Structural MRI protocols

Magnetic resonance imaging (MRI) was performed on a 3 Tesla scanner (MAGNETOM Verio, Siemens Healthcare GmbH, Erlangen, Germany) with a 32-channel head coil. Fluid-attenuated inversion recovery (FLAIR) images were acquired using the following parameters: axial acquisition orientation, 28 slices, TR=10000 ms, TE=90 ms, TI=2500 ms, FA=180°, pre-scan normalization, echo spacing=9.98 ms, bandwidth=199 Hz/pixel, FOV=220 mm, voxel size=0.9×0.9×4.0 mm3, slice order=interleaved, duration=4 min 42 s. FLAIR scans were acquired in both 2D and 3D space. Thus, we controlled this in all analyses and found no effect. The MP2RAGE sequence was acquired to assess brain structure with a voxel resolution of 1 mm (isotropic). Importantly, these T1-weighted images differ from MPRAGE T1-weighted images as they are uniform and free of other imaging properties (i.e. proton density, T2*), which can affect morphometric measurements (Babayan et al., 2019). The parameters of this sequence were the following: sagittal acquisition orientation, one 3D volume with 176 slices, TR=5000 ms, TE=2.92 ms, TI1=700 ms, TI2=2500 ms, FA1=4°, FA2=5°, pre-scan normalization, echo spacing=6.9 ms, bandwidth=240 Hz/pixel, FOV=256 mm, voxel size= 1 mm isotropic, GRAPPA acceleration factor 3, slice order=interleaved. The total acquisition time for MP2RAGE was 8 min 22 s.

#### 2.2.2. EEG protocol

An 8-min eyes-open resting state EEG was recorded with a BrainAmp MR plus amplifier in an electrically shielded and sound-attenuated EEG booth using 62-channel (61 scalp electrodes plus 1 electrode recording the VEOG below the right eye) active ActiCAP electrodes (both Brain Products GmbH, Gilching, Germany) attached according to the international standard 10–20 extended localization system, and referenced to FCz. The ground was located at the sternum, and skin-electrode impedance was kept below 5 KΩ. The amplitude resolution was set to 0.1 μV. EEG was recorded with a bandpass filter between 0.015 Hz and 1 kHz and digitized with a sampling rate of 2500 Hz. Participants were seated in front of a computer screen and asked to stay awake while fixating their eyes on a black cross presented on a white background (Babayan et al., 2019).

#### 2.2.3. Cardiometabolic measures and physical examination

A blood sample of approximately 70 ml was collected after acquiring MRI data. The blood collection utilized four distinct sampling tube types: Serum, EDTA, Citrate, and RNA. These samples were promptly dispatched on the same day to the Institute for Laboratory Medicine, Clinical Chemistry, and Molecular Diagnostics (ILM) at the Medical Faculty of Leipzig University. The selected parameters for this study included hemoglobin (HBA1c), total cholesterol, low-density lipoprotein (LDL), high-density lipoprotein (HDL), triglycerides, C-reactive protein, and creatinine.

Classical anthropometric measurements were conducted using standardized procedures by trained medical personnel. The body weight of participants, attired in clothing, was measured using an electronic scale (SECA 813, Seca Gmbh & Co KG) with a precision of 0.01 kg. Body height was measured using a stadiometer (SECA 216) with an accuracy of 0.1 cm. Waist and hip measurements were taken with an ergonometric circumference measuring tape (SECA 201), accurate to the nearest 0.1 cm. Resting blood pressure (BP) was assessed using an automatic oscillometric blood pressure monitor (OMRON M500, OMR HEM-7213-D) and a 22–42 cm arm cuff (OMRON HEM-RML30, both OMRON Medizintechnik Handelsgesellschaft mbH, Mannheim, Germany) following a seated resting period of 5 minutes, conducted before the MRI session.

### 2.3. Preprocessing and Analysis

#### 2.3.1. White matter hyperintensities

Automatic detection of white matter lesions in FLAIR images was performed with the lesion segmentation tool (LST) running on SPM12. The lesion growth algorithm (LGA), included in LST, was used (REF). The LGA calculates a lesion probability score for each voxel, generating lesion probability maps. These maps are smoothed using a Gaussian kernel with a Full Width at Half Maximum of 1 mm for voxels with a lesion probability exceeding 0.1. WMHs are commonly subdivided into periventricular and subcortical (Alber et al., 2019) each with putative neuropathological substrates related to neurovascular burden. Due to a natural volume over-representation of periventricular lesions in whole-brain volume averages, we report the total number of WMHs in the brain. The total number of WMHsdetected by the LGA algorithm was quantified per subject (Fig 1A).

**Figure 1.**
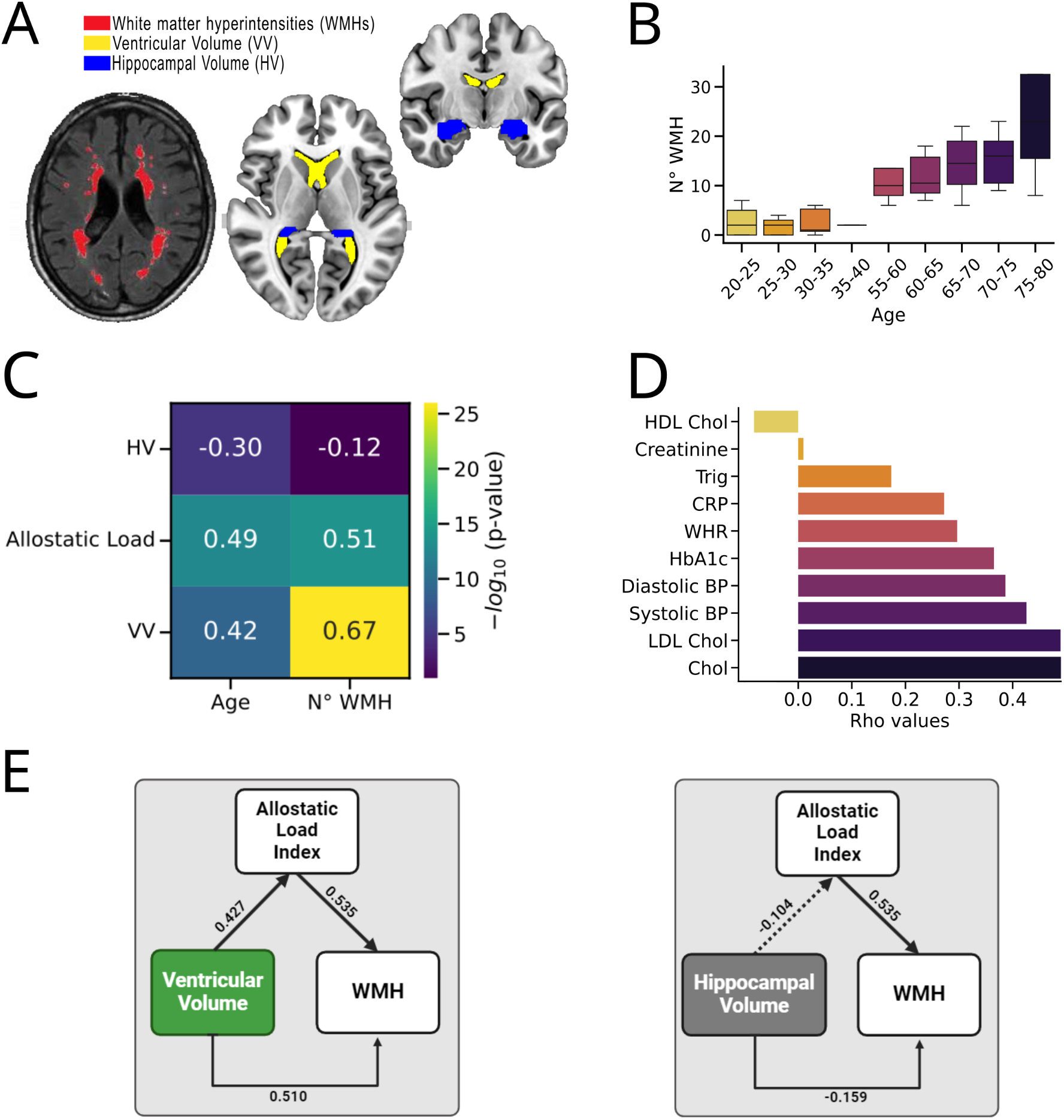
Ventricular enlargement, not hippocampal volume, is related to WMHs and allostatic load index. **A)** the axial FLAIR image on the left shows white matter hyperintensities (WMH), as red patches indicate. On the right, brain maps show anatomical localizations of bilateral ventricles (yellow) and bilateral hippocampus (blue). **B)** Bar plot showing the number of WMHs (Y axis) per age group in our sample. **C)** Correlation map showing the association of Age and Gender with Ventricle volume (VV), Allostatic load, and Hippocampal volume (HV). The figures inside each tile represent the Spearman correlation coefficients. In contrast, the coloring represents the strength in the significance of the association, with dark blue representing no significance and light to bright colors representing stronger significance. **D)** Correlations between individual biomarkers of the Allostatic load Index (Y axis) and the Number of WMH. The labels on the X-axis represent the Rho values. Also, note that separate correlations were made for each individual biomarker. **E)** Mediation analyses. The leftmost pane shows a significant mediation effect of the Allostatic load Index in the relationship between Ventricular volume and WMH. In contrast, the right pane shows no mediation of the allostatic load index in the relationship between hippocampal volume and WMH. Note that continuous and dotted lines represent significant and non-significant relationships, respectively. Figures beside each arrow indicate **β** coefficients associated with the relationship between the corresponding variables. Abbreviations: HDL: high-density lipoprotein; Chol: Cholesterol; Trig: Triglycerides; CRP: C-Reactive protein; WHR: Waist-to-hip ratio; HbA1c: Glycated hemoglobin; BP: blood pressure; LDL: low-density lipoprotein.

#### 2.3.2. MRI

The volumetric analysis of FLAIR images was conducted utilizing FastSurfer (https://deep-mi.org/research/fastsurfer/). First, FLAIR images were pre-processed using the FastSurfer pipeline, encompassing skull stripping, intensity normalization, gray and white matter segmentation, and surface reconstruction. Subsequently, the resulting gray matter was aligned to a common template space to facilitate voxel-wise analysis and corrected by the total intracranial volume of each subject (TIV). Specifically, we focused on the volume measurements of bilateral hippocampi and lateral ventricles using the Desikan-Killiany cortical atlas (Desikan et al., 2006) included in the FastSurfer software (Henschel et al., 2020)(Fig 1A).

#### 2.3.3. EEG

The processing of EEG signals was performed using NeuroDSP (Cole et al., 2019), an advanced signal processing toolbox designed for robust and comprehensive analysis of neural oscillations. Initially, raw EEG data underwent preprocessing to remove artifacts and noise, followed by segmentation into epochs of interest. Subsequently, spectral analysis was performed to extract the oscillatory features, including power spectral density and frequency bands of interest, such as alpha. Additionally, we calculated the aperiodic 1/f exponent using the FOOOF toolbox (Donoghue et al., 2020) with a fitted frequency range of 1-40 Hz.

#### 2.3.4. Allostatic load index

Here we used a composite measure of allostatic overload (Beese et al., 2022; Guidi et al., 2021; Savransky et al., 2017). Based on previous research, we included circulating indicators of inflammation (c-reactive protein), lipid profile (total cholesterol, HDL, LDL), metabolic (hip to waist ratio, hba1c), and blood pressure (systolic, diastolic blood pressure) (Mauss & Jarczok, 2021; McLoughlin et al., 2020). Despite the similarity to cardiovascular risk scores, such as the Framingham Score, ALI captures separate variance related to a global cardiometabolic state beyond predictive outcomes that do not necessarily converge (Zsoldos et al., 2018). Following previous procedures, we calculated a statistical risk threshold by distributing data into quartiles and selecting those in the highest quartile in all indicators (except HDL, for which we selected those in the lowest quartile). To calculate the composite score, a 1 was assigned for each systemic indicator whose value exceeded the risk threshold, and finally, a linear summation of all risks was calculated.

#### 2.3.5 Statistical analyses

We used Spearman Rank Correlation and partial Spearman Rank Correlation. For the correlation matrices, we report the absolute values of the 10th logarithm of the p-values to control for the magnitude difference of the significance values. All p-values were corrected by multiple comparisons with the False Discovery Rate method (Benjamini & Hochberg, 1995). Mediation analysis was applied using a bias-correct non-parametric bootstrap method with linear regressions. Correlations, partial correlations, and mediation analyses were applied using the Pingouin Python Toolbox (Vallat, 2018).

## 3 Results

To test how cardiometabolic state relates to neurovascular burden and brain structure and function, we first characterize neurovascular burden, operationally defined as the number of white matter hyperintensities (WMHs). Our analyses showed that the number of WMHs was correlated with age (Figure 1B), using gender and education as covariates (rho=0.5, p=1.04*10^−13^). Next, a composite measure of cardiometabolic state was calculated using the Allostatic Load Index (ALI, see Methods) for comparison with n° WMH. As expected, ALI was significantly correlated with age (rho=0.4, p=2.2*10^−13^) and also strongly correlated with the n° of WMHs (rho=0.512, p=1.7*10^−14^), even when controlled by age, gender, and education (rho=0.31, p=0.000017; Figure 1C).

To explore how age-related neuroimaging biomarkers are affected by systemic cardiometabolic state, we calculated how bilateral hippocampal and lateral ventricular volume was associated with WMHs and with allostatic load index. Both ventricular volume (VV) and hippocampal volume (HV) were significantly correlated with WMHs (VV: rho=0.7, p-value=1.7^-31^; HV: rho=0.2, p-value=0.001). However, only VV was correlated with n° of WMHs after controlling for age, gender, and education (VV: rho=0.59, p-value=7.7^-20^; HV: rho=-0.02, p-value=0.71). Consistently, we found that only VV correlated with ALI (VV: rho=0.25, p-value=0.0003; HV: rho=-0.03, p-value=0.62; Figure 1C). Next, we tested if ALI mediated the relation between these brain volumes and WMHs, and found that only VV was significantly mediated by ALI, with a significant indirect (mediated) effect (β = 0.135, p = 0.00, C.I = 0.082 - 0.216). Path a (i.e., VV on ALI) (β = 0.427, p = 4.1747^-10^) and path b (i.e., ALI on WMHs) (β = 0.535, p = 5.78891^-16^) were both significant (Figure 1E). These results suggest that VV is related to WMHs mediated by cardiometabolic state but not HV, possibly pointing to different biological pathways across aging.

Next, we explored how WMHs and ALI are related to brain function, analyzing canonical age-related EEG measures, such as the power of alpha (Babiloni et al., 2006; Klimesch, 1999) and the 1/f exponent (Donoghue et al., 2020; Voytek et al., 2015). We found that both alpha and the spectral exponent were significantly related to age (alpha: rho=0.23, p=0.001; 1/f exponent: rho=-0.6, p=2.9e-21), however only the 1/f exponent correlated with WMHs (alpha: rho=0.11, p=0.1; 1/f exponent: rho=-0.4, p=8.2e-09). Consistently, we found that only the 1/f exponent was related to ALI (alpha: rho=-0.13, p=0.056; 1/f exponent: rho=-0.39, p=1.2e-08). To test if the structural aging pathway of VV and HV were related to brain function, we correlated both structural measures with alpha power and 1/f exponent. We found that both were significantly correlated with 1/f exponent and not alpha (alpha∼VV: rho=-0.136, p=0.055; alpha∼HV: rho=0.13, p=0.06; 1/f exponent∼VV: rho=-0.34, p=6.6e-07; 1/f exponent∼HV: rho=0.33, p=1.0e-06). Interestingly, we found that VV-related WMHs was associated to 1/f exponent with a complete mediation of ALI [(β = -0.156, p = 0.00, C.I = -0.238 - -0.088), path a (VV-related WMH on ALI) (β = 0.427, p = 4.1747^-10^); path b (ALI on 1/f exponent) (β = -0.425, p = 4.89391^-10^)]. However, that was not the case for HV-related WMHs, where there was a non-significant mediation of ALI (β = -0.041, p = 0.146, C.I = -0.103 - -0.103) and non-significant path a (HV-related WMH on AL) (β = 0.104, p = 0.146) (Figure 2D). Our results show that the cardiometabolic state is related to specific structural and functional pathways across aging, specifically characterized by neurovascular burden, ventricular enlargement, and EEG spectral exponent.

**Figure 2.**
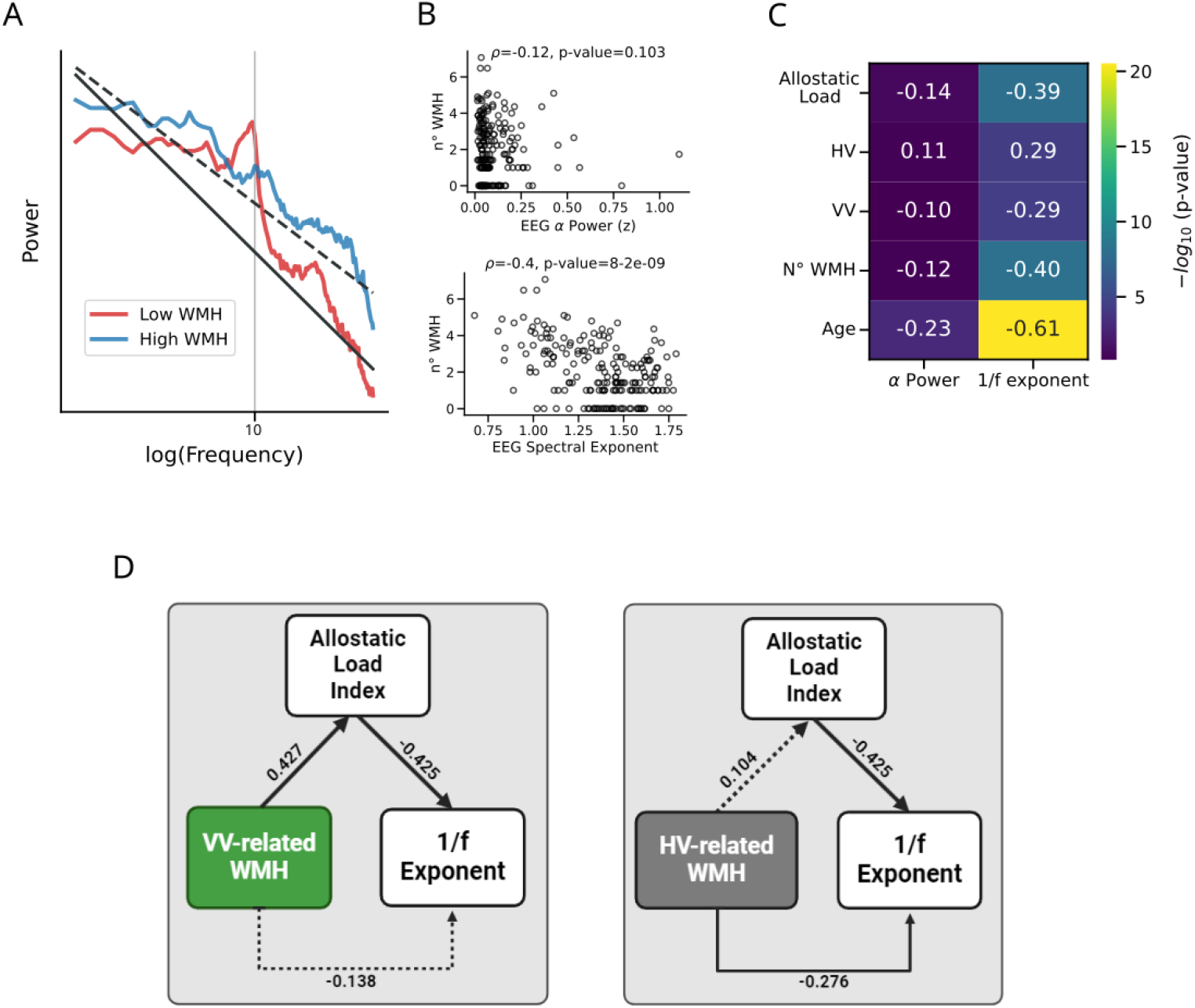
Electrophysiological correlates of cardiometabolic state, neurovascular-structural burden. **A)** Aperiodic EEG activity between low and high WMHsburden, as indicated by red and blue. **B)** The top plot shows a non-significant correlation (Spearman) between WMHsand total alpha power. The bottom plot shows the positive association between WMHsand spectral exponent (1/f). **C)** Correlation map showing the association of Total alpha power and spectral exponent with age, number of WMH, Ventricular volume (VV), hippocampal volume (HV), and allostatic load. The figures inside each tile represent the Rho values. In contrast, the coloring represents the strength in the significance of the association, with dark blue tiles representing non-significance and light to bright colors representing stronger significance. **D)** Mediation analysis: the left figure shows a complete mediation of the allostatic load index in the relationship between ventricle-related WMHsand 1/f exponent. The right figure shows a non-significant mediation of the allostatic load index on the relationship between hippocampal volume-related WMHsand 1/f exponent. Note that continuous and dotted lines represent significant and non-significant relationships, respectively. Figures beside each arrow indicate **β** coefficients associated with the relationship between the corresponding variables.

## 4. -Discussion

Across age, systems-level energy supply becomes continuously impaired from its demand. This divergence has been related to cardiac and metabolic risk factors that shape brain health. Here we leverage a composite allostatic measure to study how systemic cardiometabolic state links with WMHs, and age-related structural and functional brain measures. We define WMHs as neurovascular burden considering its strong relation with cardiovascular risk factors (Claassen et al., 2021; Moroni et al., 2018; Raz et al., 2012; Zhao et al., 2019) and cerebrovascular disease (Rastogi et al., 2021; Vemuri et al., 2021). Here, we found that WMHs are closely related to ventricular enlargement and the 1/f exponent of the EEG functional activity, mediated by the cardiometabolic allostatic state, thus expanding our previous knowledge on the functional and structural nature of WMHs in the human brain. Consistent with prior work showing that WMHs are related to local brain metabolism (Brier et al., 2022; Jiaerken et al., 2019), we show that global measures of cardiometabolic state affect both structural and functional brain properties related to the emergence of neurovascular burden.

Ventricular and hippocampal volume -the most widely used structural biomarker of aging (Izzo et al., 2020; Uysal & Ozturk, 2020; Klistorner et al., 2022)- are generally inversely correlated and change across the lifespan (Bethlehem et al., 2022). Both volumetric measures have been linked to age-related pathologies, such as Alzheimer’s disease (Jack Jr et al., 2000; Medel et al., 2024; Nestor et al., 2008). However, they have been postulated to have alternative biological pathways. The loss of hippocampal volume is related to structural changes in neurodegenerative processes associated with neuronal loss. In contrast, ventricular enlargement is associated with cerebrospinal fluid inflammatory processes, which can even be reversible (Millward et al., 2020; Zahr et al., 2013), suggesting that although both volumetric measures co-occur in aging, they do not necessarily share the same mechanisms. Our results highlight the cardiometabolic pathway of ventricular enlargement and its association with neurovascular burden as a putative neuroinflammatory pathway of brain aging beyond hippocampal and gray matter atrophy (Supplementary Figure 1).

Neural function is also highly sensitive to age. Using two canonical age-related EEG proxies, we found that alpha power and the 1/f exponent of the spectrum both correlated with age (Figure 1B). Interestingly, only the aperiodic exponent was correlated with allostasis and WMHs. Moreover, we found that an increase in WMHs related to ventricular enlargement and not hippocampal atrophy was related to 1/f exponent with absolute mediation of the allostatic load. This result suggests that the cardiometabolic state links structural imprinting of neuroinflammatory pathways seen in WMHs and ventricular enlargement with EEG aperiodic activity. Although it has been shown that the 1/f exponent is related to neuronal circuit properties such as the balance between excitation and inhibition (Gao et al., 2017; Medel et al., 2023; Trakoshis et al., 2020), recent evidence links this global electrophysiological signature with physiological signals, such as oxygen availability (Coronel-Oliveros et al., 2024), respiration (Kluger et al., 2023) and cardiac measures (Schmidt et al., 2022). Our work provides evidence that this relation does not suggest an artefactual origin but rather a natural consequence of the close relation between electrophysiological function, measured by 1/f exponent background signal, and the energetic requirements of neural circuits.

## Limitations

Here we draw a sample from a specific region that may not be representative of other geographic or ethnic populations, as cultural lifestyle and environmental stressors can influence both cardiometabolic health, brain structure and function, as well as increased allostatic load (Migeot et al., 2024). Thus, further studies can use data considering the recent efforts to increase diversity in structural and functional age-related brain measures (Hernandez et al., 2024; Prado et al., 2023), Migeot et al., 2024). Additionally, our analyses were done in a cross-sectional sample which makes causal relationships elusive, requiring longitudinal data to explore the temporal relationship between cardiometabolic state, WMHs, with brain structure and function. Despite this potential limitation, previous evidence has shown that local inflammation tended to generate WMHs after 1 year of monitoring, suggesting that cardiometabolic and inflammatory mechanisms are indeed causally related to WMHs (Tozer et al., 2023). These mechanisms have been widely implicated in diverse age-related pathologies, such as Alzheimer’s Disease and other neurodegenerative disease (Luo et al., 2023), suggesting that cardiometabolic allostatic state might be underlying the alteration of structural and functional measures of the pathological brain (Adedeji et al., 2023; Birba et al., 2022; Ibanez, 2023). Moreover, lifestyle, exposure to stressful events, poor nutrition, or low access to health care have all been related to social determinants of health associated with cardiometabolic state (Alvarez et al., 2022; Breitenbach et al., 2021; Evans et al., 2021; Lucente & Guidi, 2023; Ribeiro et al., 2019). Our work paves the way for future characterization of cardiometabolic aging pathways mechanistically associating brain structure and function, putatively affected in populations with high social determinants of health risk (Prado et al., 2023). This offers a new window to understanding the general mechanisms of aging and broadening the diagnostic and therapeutic tools related to healthy and pathological aging.

## Conclusions

In this study, we investigated the connections between cardiometabolic health and brain structure and function across age. Our findings underscore the critical impact of cardiometabolic health on brain integrity and dynamics, suggesting neuroinflammatory mechanisms underlying WMHs, ventricle enlargement, and increased aperiodic exponent across aging. The results also highlight the importance of preventing cardiometabolic risk factors to maintain healthy brain aging and could inform strategies aimed at preserving brain health through targeted interventions addressing cardiometabolic risk factors.

## Author’s contribution

Conceptualization, Data Curation, Writing - Original Draft: DFO, ACL, AI and VM. Methodology, Investigation, Software, Formal Analysis: CD, JMS, DH, ET, CC, DFO, ACL, AI and VM. Writing - Review and Editing: CD, JMS, DH, ET, CC.

## Notes

### Competing Interest Statement

The authors have declared no competing interest.

