## Supplemental Figure 1 for "Cardiometabolic state links neurovascular burden with brain structure and function across age: evidence from EEG and MRI"

**Supplementary 1**

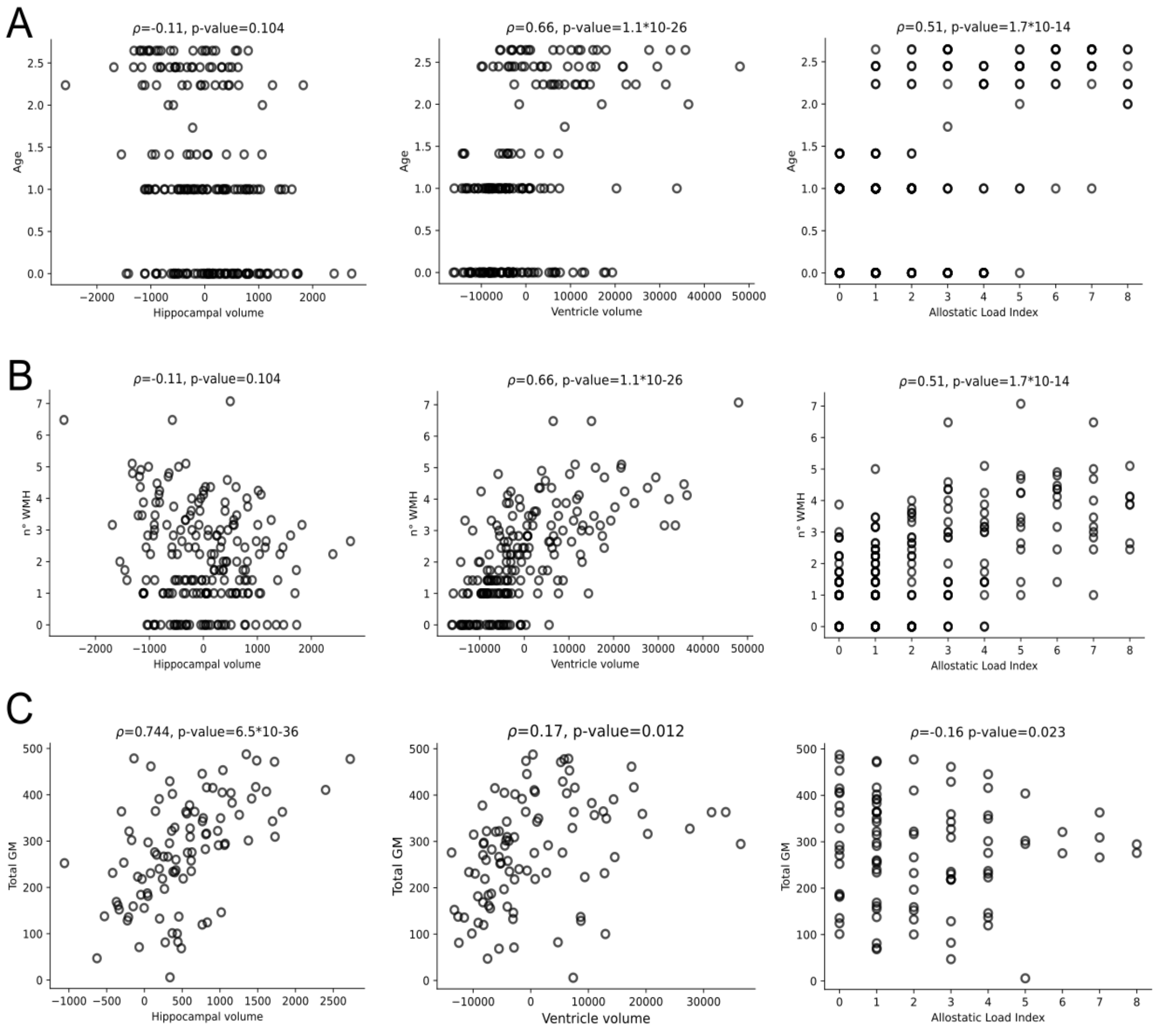


Correlation analyses examining the association between **A)** Age and Hippocampal volume, Ventricle volume, and Allostatic load index; **B)** White matter hyperintensities and Hippocampal volume, Ventricle volume, and Allostatic load index; **C)** Total Gray Matter (GM) and Hippocampal volume, ventricle volume, and Allostatic load index.
